# Comparative transcriptomics provides insight into molecular mechanisms of zinc tolerance in the ectomycorrhizal fungus *Suillus luteus*

**DOI:** 10.1101/2023.12.08.570832

**Authors:** Alexander Smith, Janne Swinnen, Karl Jonckheere, Anna Bazzicalupo, Hui-Ling Liao, Greg Ragland, Jan Colpaert, Anna Lipzen, Sravanthi Tejomurthula, Kerrie Barry, Igor Grigoriev, Joske Ruytinx, Sara Branco

**Affiliations:** Department of Integrative Biology, University of Colorado Denver, Denver CO, USA; Research Groups Microbiology and Plant Genetics, Vrije Universiteit Brussel; Comparative Fungal Biology, Royal Botanic Gardens, Kew, Richmond, United Kingdom; Soil, Water and Ecosystem Sciences Department, University of Florida, Gainesville, Florida, USA; North Florida Research and Education Center, The University of Florida, Quincy, Florida, USA; Hasselt University; DOE Joint Genome Institute, Lawrence Berkeley National Laboratory, Berkeley, CA, USA; Department of Plant and Microbial Biology, University of California Berkeley, Berkeley, California, USA

## Abstract

Zinc is a major soil contaminant and high zinc levels can disrupt growth, survival, and reproduction of fungi. Some fungal species have evolved zinc tolerance through cell processes mitigating zinc toxicity, though the genes and detailed mechanisms underlying fungal zinc tolerance remain unexplored. To fill this gap in knowledge, we investigated the gene expression of zinc tolerance in the mycorrhizal fungus *Suillus luteus*. We found that zinc tolerance in this species is both a constitutive and environmentally dependent trait. Highly differentially expressed genes were predicted to be involved in transmembrane transport, metal chelation, oxidoreductase activity, and signal transduction. Some of these genes were previously reported as candidates for *S. luteus* zinc tolerance, while others are reported here for the first time. Overall, we found *S. luteus* zinc tolerance is associated with differences in expression of genes involved in metal exclusion and immobilization, as well as recognition and mitigation of metal-induced oxidative stress. Our results contribute to understanding the mechanisms of fungal metal tolerance and pave the way for further research on the role of metal tolerance in mycorrhizal associations.

## Introduction

Zinc (Zn) is an essential micronutrient (Robinson et al., 2021) critical to biological functions such as the regulation of nucleic acids, enzyme activation, and the synthesis of proteins, carbohydrates, and lipids (Sharma et al., 2013). However, at high concentrations Zn is toxic and can lead to death. Alarmingly, increased anthropogenic activity has caused Zn contamination to become one of the most prevalent and detrimental forms of pollution in soil environments (Guo et al., 2014, Smith et al., 2016, Ramrakhiani et al., 2017, Op de Beeck et al., 2015). High Zn environments impose stressful conditions such as altered soil chemical properties and lower nutrient bioavailability that can lead to the disruption of cellular and organismal processes responsible for metabolism, growth, and survival (Påhlsson, 1989, Priyadarshini et al., 2021, Wang et al., 2006). Despite these harsh conditions, some species tolerate high Zn levels and even thrive in Zn-rich environments (Miransari, 2011). The exact mechanisms of Zn tolerance in mycorrhizal fungi are however still unclear.

There has been increasing interest in understanding how fungi interact with Zn (Branco et al., 2022). Fungi have evolved homeostatic mechanisms to maintain the Zn levels required in cell processes such as transcription, protein folding, and hyphal growth (Feldmann, 2012). However, excess Zn negatively impacts fungal cell function by interrupting cell membrane synthesis (Galván Márquez et al., 2018), increasing oxidative stress that can trigger programmed cell death (Zadrąg-Tęcza et al., 2018), and decreasing growth by disrupting hyphal extension and altering hyphal morphology (Lanfranco et al., 2010, Priyadarshini et al., 2021). Furthermore, high Zn levels can reduce spore germination (Pawlowska and Charvat, 2004), lower fungal survival (Op De Beeck et al., 2015, Priyadarshini et al., 2021), and decrease fungal community species richness (Faggioli et al., 2019).

Even though high Zn concentrations are sub-optimal, some fungi evolved high Zn tolerance and can persist in Zn contaminated environments (Ezzouhri et al., 2009, Anahid et al., 2011, Op De Beeck et al., 2015, Robinson et al., 2021). Fungal Zn tolerance relies on regulation of Zn transport, sequestration, and immobilization, as well as the production of antioxidants that mitigate toxicity (Bellion et al., 2006). Zn import is controlled by metal transmembrane transporters and the reduction of bioavailable forms of extracellular Zn. For example, in the ericoid mycorrhizal fungus *Oidiodendron maius*Zn tolerance relies on the plasma membrane Zn transport protein OmFET and isolates collected from Zn-polluted soils showed a much lower ability to solubilize inorganic Zn (Khouja et al., 2013, Martino et al., 2003). Additionally, the ectomycorrhizal *Hebeloma cylindrosporum* sequesters cytosolic Zn into intracellular vesicles through a transport protein localized to the endoplasmic reticulum (Blaudez and Chalot, 2011). Fungi can also reduce the amount of excess cytosolic Zn through metallothioneins, metal complexing proteins that immobilize micronutrient metals (Howe et al., 1997). Finally, antioxidant homeostatic mechanisms produce enzymes that break down reactive oxygen species (ROS), neutralizing oxidative stress toxicity resulting from high intracellular metal content (Colpaert, 2008, Bolann and Ulvik, 1997, Teng et al., 2017).

Studies in *Suillus luteus* have substantially contributed to unveiling the mechanisms of fungal Zn tolerance. *S. luteus* is a widespread temperate ectomycorrhizal fungus that associates with pine trees, maintaining a mutualistic relationship where it provides mineral nutrients and receives carbohydrates in return (Lofgren et al., in review). *S. luteus* is an important pioneer species and dominates fungal communities in metal-contaminated sites (Colpaert and van Assche, 1987, Ruytinx et al., 2011, Op De Beeck et al., 2015). This species displays Zn tolerance in areas around decommissioned Zn smelters in Belgium, where some isolates can survive at high Zn soil concentrations (Colpaert and van Assche, 1987, Colpaert et al., 2004). Notably, Belgian *S. luteus* Zn tolerant isolates accumulate Zn in their tissue at a much slower rate and can withstand much higher concentrations as compared to sensitive isolates (Colpaert et al., 2005). Genetic studies on these isolates identified four transmembrane transporter genes involved in cellular Zn homeostasis and putatively involved in tolerance. *SlZnT1* and *SlZnT2* are cation diffusion facilitator (CDF) family Zn transporters involved in vacuolar transport, suggesting that intracellular metal sequestration plays an important role in *S. luteus* Zn tolerance (Ruytinx et al., 2017). In addition, the plasma membrane Zrt/IrT-like protein (ZIP) transporters *SlZRT1* and *SlZRT2* showed significant downregulation in high Zn environments, indicating they also play a key role in maintaining Zn homeostasis (Coninx et al., 2017, 2019). Furthermore, genome scans of the same Belgian *S. luteus* isolates revealed the absence of population structure between Zn contaminated and non-contaminated soils, and that metal tolerance in this species is polygenic (Bazzicalupo et al., 2020). Genetically differentiated loci between isolates from contaminated and non-contaminated sites included transmembrane transporters, chelators, and antioxidants, namely proteins involved in Zn import and vacuolar sequestration, intracellular and extracellular Zn immobilization, and ROS detoxification. Interestingly, *SlZnT1, SlZnT2, SlZRT1* and *SlZRT2* did not show high genetic divergence (Fst) within coding regions across isolates from contaminated and non-contaminated sites isolates (Bazzicalupo et al., 2020). These results suggested that Zn transporter genes may instead confer Zn tolerance through differential expression.

Here, we quantify gene expression differences in previously studied Belgian *S. luteus* isolates from contaminated and non-contaminated soils to unveil the mechanisms of fungal Zn tolerance. We hypothesized that isolates originating from contaminated soils would be Zn tolerant and isolates from non-contaminated soils would be sensitive to high Zn concentrations. Because these isolates belong to the same population and are genetically very similar (Bazzicalupo et al, 2020), we expected to find significantly different transcriptomic profiles only across high and low Zn treatments, including in the candidate genes highlighted in previous experiments. We also predicted that tolerant and sensitive isolates would show distinct responses to the Zn treatment, with differentially expressed genes contributing to Zn tolerance pathways and mechanisms in *S. luteus*. We found that the level of soil contamination was positively associated with isolate Zn tolerance and that transcriptomic differences between tolerant and sensitive isolates were both constitutive and Zn induced. We also document *S. luteus* putative Zn tolerance mechanisms, including metal exclusion and immobilization, as well as recognition and mitigation of metal-induced oxidative stress. Our analyses provide a transcriptome-wide exploration of potential ectomycorrhizal fungal Zn tolerance mechanisms and highlight many genes likely important to Zn tolerance in *S. luteus*.

## Experimental Procedures

### Sampling sites and fungal culture information

To investigate the effect of Zn on *S. luteus* gene expression, we studied ten isolates from Belgium, five collected from one soil contaminated site (Lommel) and five collected from one non-contaminated site (Paal) (Colpaert et al., 2004, Supplemental Table 1). All ten isolates show little overall genetic differentiation and belong to a single population (Bazzicalupo et al, 2020). Both contaminated and non-contaminated sites are dominated by Scots pine (*Pinus sylvestris*) and the contaminated site displays high levels of Zn, Pb, Cd, Cu, and As resulting from decommissioned zinc smelters (Op De Beeck et al., 2015; Sonke, Hoogewerff, van der Laan, & Vangronsveld, 2002). We obtained *S. luteus* pure cultures from fruitbodies and maintained them in solid modified Fries medium (Supplemental Table 2; Colpaert 2000). For more information on collection methods and site descriptions consult Bazzicalupo et al. (2020).

### Testing *S. luteus* Zn tolerance

To quantify *S. luteus* Zn tolerance and whether it correlates with soil contamination, we calculated EC_50_ values (amount of Zn that inhibits isolate growth by 50%) for each of the ten isolates. We grew the isolates at 23°C in darkness on cellophane-covered solid modified Fries medium supplemented with different concentrations of ZnSO_4_·7H_2_O (0, 100, 200, 400, 800, and 1000 ppm). We performed four replicates per isolate for each condition. After 14 days of growth, we collected the mycelium from the plates and stored it at −80°C. To determine EC_50_ values per isolate, we followed the protocol in Ritz et al (2015). Briefly, we obtained isolate dry weight after lyophilization and used it to construct dose-response curves through non-linear regression with a four-parameter log-logistic model in R version 3.5.1 (R Core Team, 2018). We used EC_50_ values to place isolates into tolerance categories (tolerant, sensitive) irrespective of collection site. We considered isolates Zn tolerant if EC_50_ > 130 ppm and sensitive if EC_50_ < 130 ppm as this was the amount of Zn added in our transcriptomic experiment.

### RNA Extraction and Sequencing

To measure Zn-induced differential gene expression across the six Zn tolerant and four Zn sensitive *S. luteus*, we grew the isolates on both control and zinc-supplemented (130 ppm Zn) modified solid Fries medium covered with cellophane, in triplicate, at 23°C, and in darkness (n = 10 isolates x 2 treatments x 3 replicates = 60). We selected 130 ppm Zn because this concentration is sublethal to sensitive isolates. Once the isolate on the control plate reached a diameter of 3 cm (10-14 days, depending on the isolate), the mycelium for both control and corresponding Zn supplemented plates was collected and stored at −80°C awaiting RNA extraction.

We extracted total RNA using the RNeasy Plant Mini Kit (Qiagen, France) on mycelia ground in liquid nitrogen using mortar and pestle. We measured the RNA concentration and determined the A260/230 and A260/280 ratios using a Nanodrop One (Thermo Fisher). Samples that did not meet purity criteria (A260/230 and A260/280 > 1.8) were discarded. RNA integrity was confirmed using a Bioanalyzer with the RNA 6000 Nano Kit (Agilent). Two sample replicates (one P1 and one P13, both grown in 130 ppm Zn) did not meet our quality criteria and were not incorporated into subsequent analyses. RNA was sequenced by the Joint Genome Institute (JGI) on an Illumina NovaSeq 6000 S4 (2 x 151 bp) and run through JGI’s quality and quantity QC tests (https://jgi.doe.gov/user-programs/pmo-overview/project-materials-submission-overview/). Three sample replicates did not meet sequence quality standards (two from isolate P2 and one from isolate PD13 all in 130 ppm Zn) and were discarded, so that our final number of samples for gene expression analysis was reduced to 55.

### Bioinformatics Pipeline

The RNA sequence analyses involved four steps: read quality control, alignment to reference genome and raw gene counts, differential gene expression analyses, and gene ontology analyses:

#### Read quality control and preprocessing

We used BBDuk (Bushnell, 2016) to filter and trim raw reads. We used kmer matching (kmer=25) to remove Illumina adapters, sequencing artifacts, RNA spike-in reads, PhiX reads, and reads containing any ‘N’s. We used the phred trimming method to trim the read ends, set to Q6. Finally, we removed reads shorter than 25 bases or 1/3 the length of the original read.

#### Alignment to reference genome and raw gene counts

We used HISAT2 (Kim et al., 2015) to align reads to the *S. luteus* reference genome, calculate gene counts, and extract gene annotations. The reference genome (*S. luteus* isolate UH-Slu-Lm8-n1) originated from the same Belgian population as the isolates used in our study (JGI Project ID: 1006871; Kohler et al., 2015). To calculate raw gene counts, we used the program featureCounts (Liao et al., 2014) and the gff3 annotation file from the reference genome. Specifically, we only used primary hits on the reverse strand for the final gene counts (-s 2 -p -- primary options).

#### Differential Gene Expression Analysis

To characterize the differences in transcriptomic response to Zn between tolerant and sensitive isolates, we fit a generalized (negative binomial) linear model using the R package DESeq2 (Love et al., 2014) including parameters for Zn treatment, tolerance group, and the interaction between these two effects. We calculated a Benjamini-Hochberg adjusted p-value (FDR) for each parameter using Wald tests and used a threshold of FDR < 0.05 as adequate evidence that a parameter improved the fit of the model. We then used fitted models and associated FDRs to identify three groups of interest: 1) transcripts that were differentially expressed in response to the Zn treatment irrespective of the sensitivity of the isolate (Zn treatment FDR < 0.05; interaction FDR > 0.05), 2) transcripts that were differentially expressed between tolerant and sensitive isolates irrespective of Zn treatment (tolerance group FDR < 0.05, interaction FDR > 0.05), and 3) transcripts whose differential expression in response to Zn treatment differed between tolerant and sensitive isolates (interaction FDR < 0.05). These categories allow us to interpret the direction and magnitude of differential expression in a biologically relevant manner. To visualize clustering of samples based on overall transcript expression in the model, we used the *plotPCA* function of DESeq2 to run a principal component analysis (PCA). To test for differences between groups in the model, we ran a PERMANOVA using the Jaccard index to calculate pairwise distances and 9999 permutations (vegan package: Oksanen et al., 2008). We then calculated adjusted p-values using the Benjamini-Hochberg method.

#### Gene Ontology (GO) Enrichment Analysis

To identify enriched (overrepresented) gene annotation categories in the three groups of transcripts, we implemented an enrichment analysis using ClueGO (v2.5.9) (Bindea et al., 2009), a Cytoscape (Shannon et al., 2003) application. This analysis used the Gene Ontology (GO) annotation database to extract gene annotation categories that were enriched in each of the transcript groups as compared to the whole genome. We investigated GO annotation terms from all three overarching categories (biological process, molecular function, and cellular component) and used modified settings for both the GO Tree Interval (min = 3; max = 15) and GO Term/Pathway Selection (min # genes = 1; % genes = 4) to include the most GO terms possible. We used two-sided hypergeometric tests with a Benjamini-Hochberg (FDR) correction to weight evidence for enrichment, reporting only GO terms with an FDR less than or equal to 0.05. Additionally, to identify highly differentially expressed genes, we restricted our analysis to transcripts only with an absolute log fold change value greater than 1.

## Results

### *Suillus luteus* Zn Tolerance Correlates with Soil Contamination

We found *S. luteus* Zn tolerance is highly associated with soil contamination (ANOVA: F^1,8^ = 10.38, p = 1.22e^-02^). Isolates collected from the contaminated site were more Zn tolerant than isolates collected from the non-contaminated site, with the most tolerant isolate having an EC_50_ concentration 15 times larger than the most sensitive isolate (Figure 1). Interestingly, one isolate from the non-contaminated site was more Zn tolerant than one of the isolates from the contaminated site. The four sensitive isolates had EC_50_ values considerably lower than the Zn concentration used in the experiment (130 ppm). Conversely, the most Zn tolerant isolate’s EC_50_ concentration was 7 times greater than the amount of added Zn and may not have been as affected by the Zn addition. Based on these EC_50_ values, we divided the ten isolates into Zn tolerant and sensitive groups (Figure 1) as detailed in the methods.

**Figure 1.**
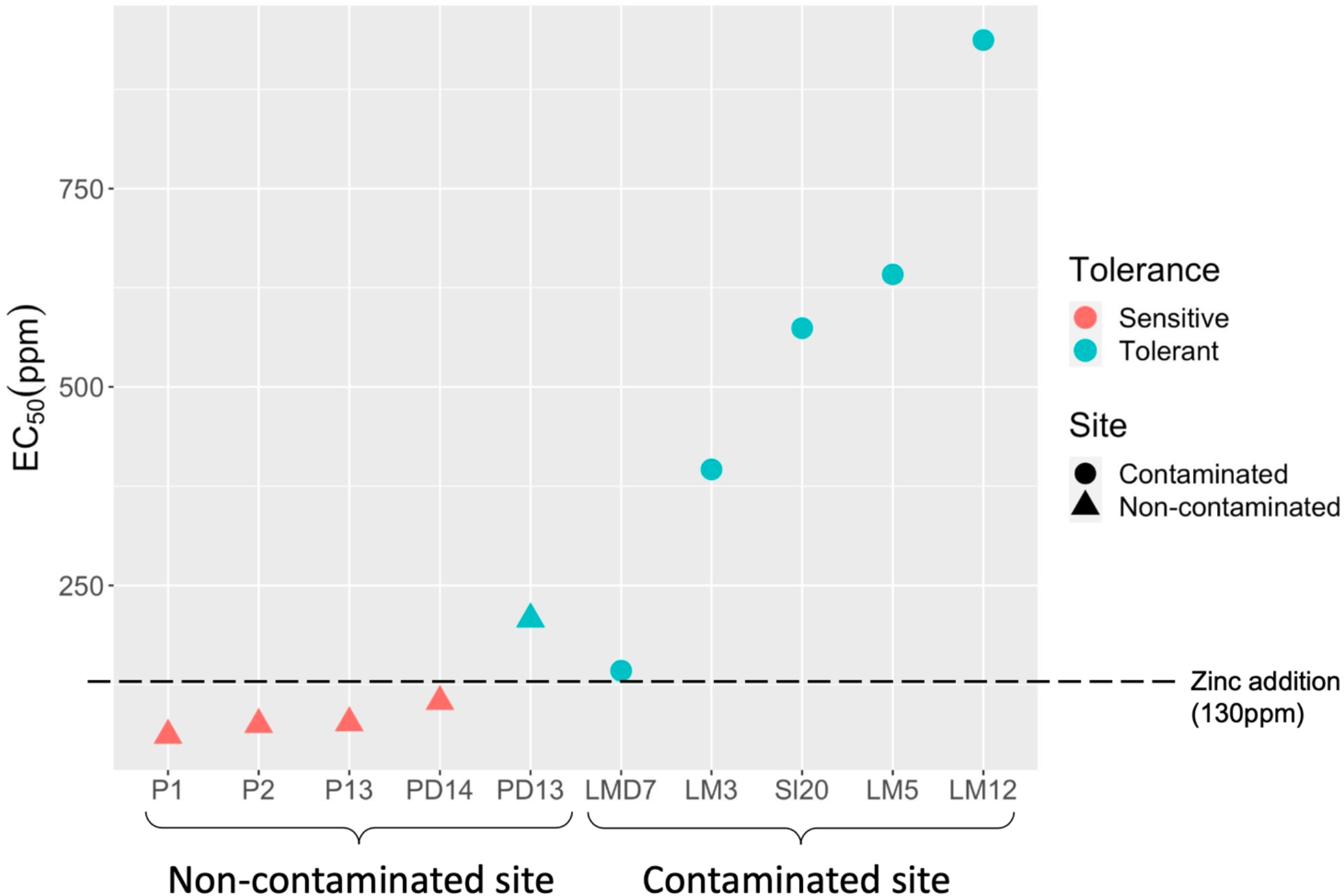
Zn Tolerance correlates with soil contamination. EC50 values for the studied *S. luteus* isolates, grouped by collection site. The shape of the symbols denotes collection site (triangles = non-contaminated site, circles = contaminated site), and the color of the symbols denotes tolerance group (red = sensitive, blue = tolerant). Dashed line indicates 130 ppm Zn, the amount of Zn added to the Zn-treated plates.

### Zn Tolerant *S. luteus* Isolates Have Higher Transcriptomic Variation Compared to Sensitive Isolates

The principal component analysis showed that *S. luteus* isolates displayed significant transcriptomic variation across Zn tolerance groups (PERMANOVA: F_1,51_ = 4.98, p-adj = 3.00e^-04^) and Zn treatment (PERMANOVA: F_1,51_ = 2.71, p-adj = 3.45e^-03^; Figure 2). The Zn tolerant isolates showed much higher transcriptomic variation compared to the sensitive isolates (Figure 2). We did not find a significant tolerance group-dependent Zn effect (interaction effect) (PERMANOVA: F_1,51_ = 1.06, p-adj = 0.366), indicating that tolerant and sensitive isolates do not show significantly different overall transcriptomic variation under Zn exposure.

**Figure 2.**
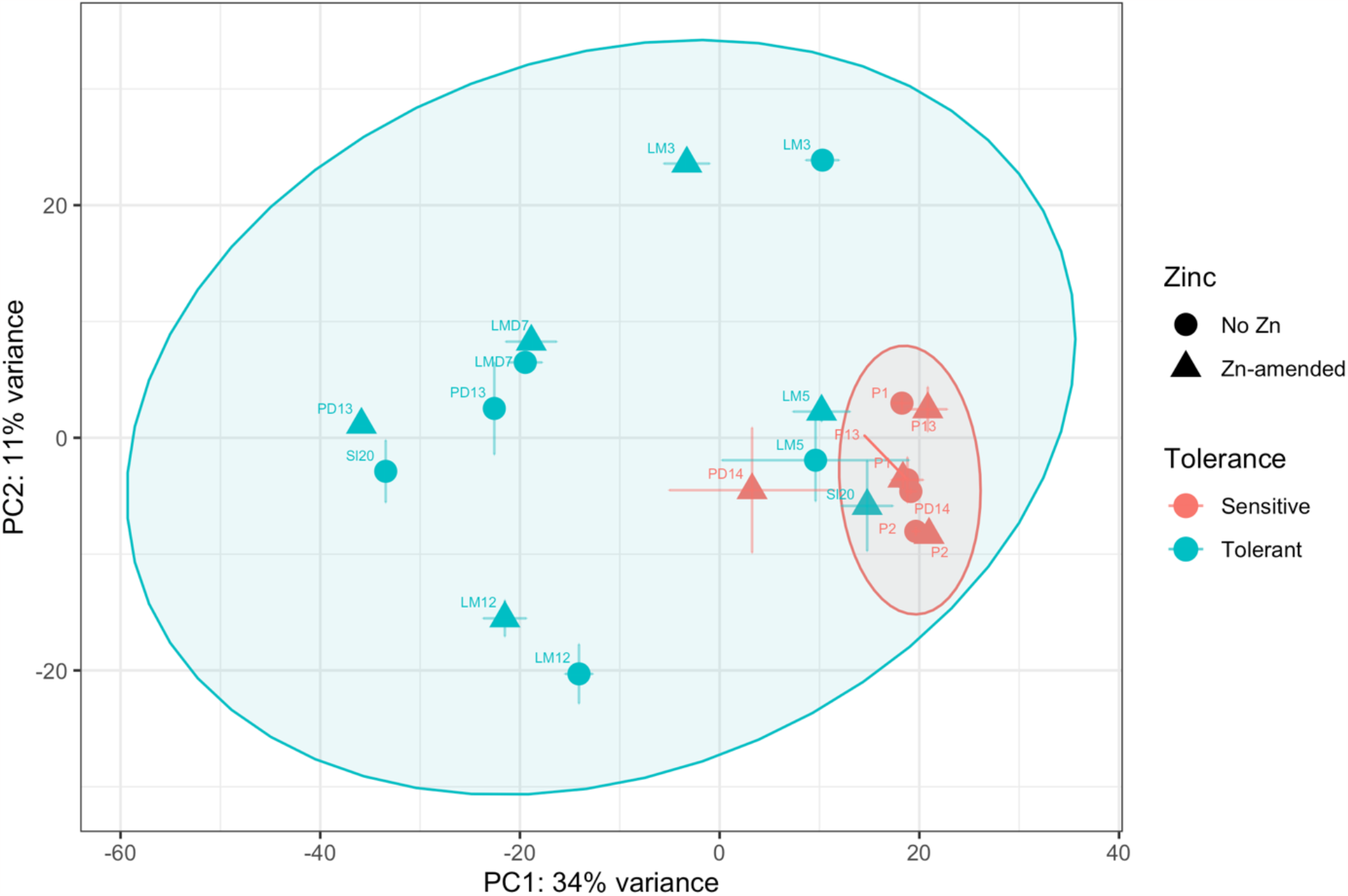
Tolerant isolates show higher transcriptomic variation compared to sensitive isolates. Principal Component Analysis plot of complete transcriptomes of the Zn sensitive and tolerant *S. luteus* isolates. Symbols represent the average across isolate replicates and error bars represent the standard error between replicates. Colors denote tolerance grouping (red = sensitive, blue = tolerant) and the symbol shape denotes Zn treatment (circle = no zinc, triangle = zinc amended). We include isolate identifiers next to each symbol. Ellipses denote 95% confidence intervals for each of the tolerance groups (red = sensitive, blue = tolerant).

### Isolate Zn Tolerance Level, not Zn Treatment, Shows Largest Number of Differentially Expressed Genes

The differential gene expression analysis showed that many more genes were differentially expressed between isolate Zn tolerance groups than between Zn treatment. Specifically, we found that over 80 times as many genes were significantly differentially expressed between tolerant and sensitive isolates as compared to Zn treatment (Figure 3). In addition, we detected 4098 genes for which tolerant and sensitive isolates responded to Zn differently (interaction effect).

**Figure 3.**
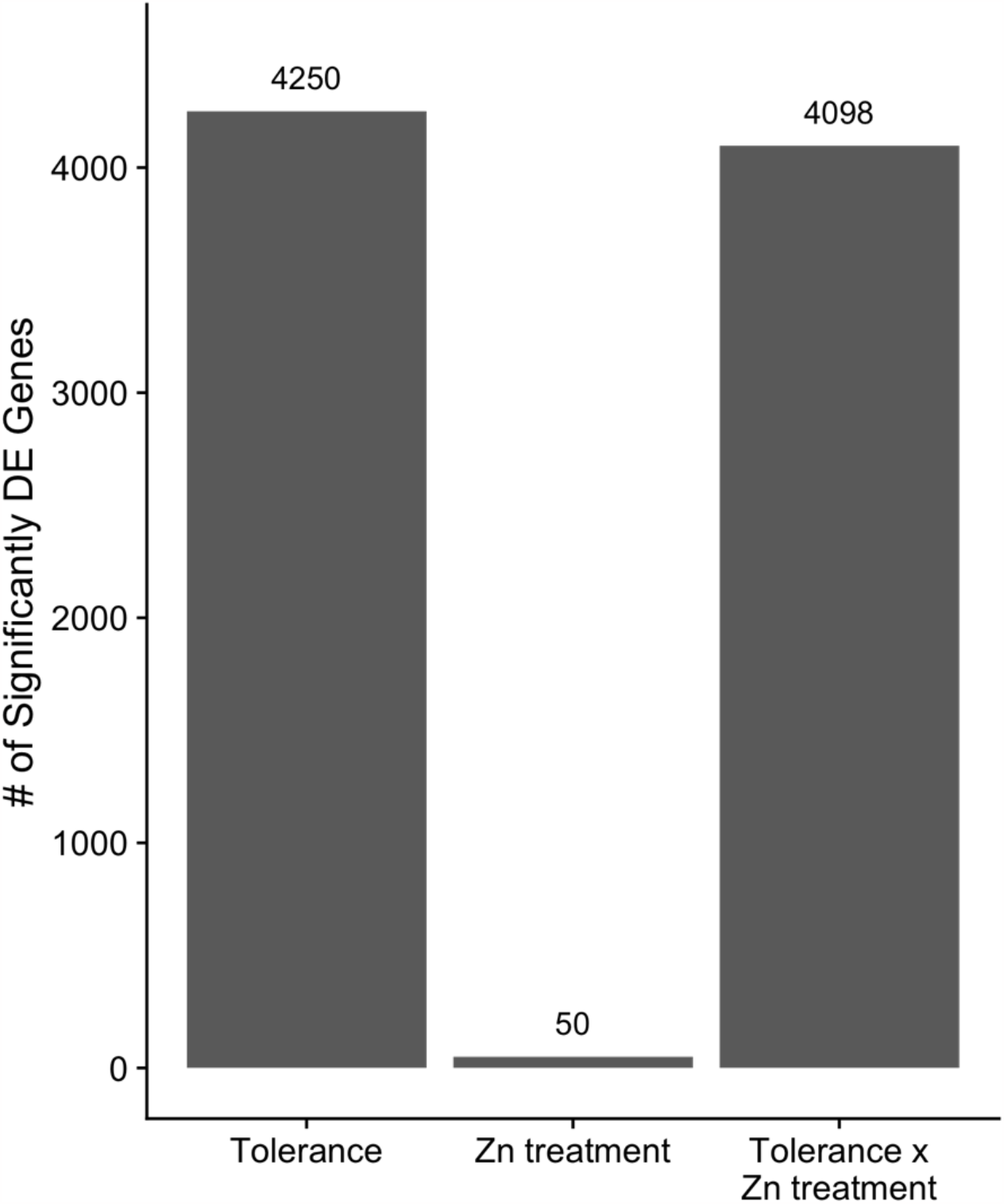
Zn tolerance affects gene expression more than Zn treatment. Barplot displaying the number of significantly differentially expressed (DE) genes in each of the three model comparisons. Tolerance = tolerance groups (tolerant vs sensitive), Zn treatment = control vs Zn-amended, Tolerance x Zn treatment = interaction effect. The numbers on top of bars represent the number of significantly DE genes. DE genes that had a significant interaction effect were removed from the DE gene counts for the main effects (Tolerance and Zn treatment).

### Enrichment of Transmembrane Transport, Oxidoreductase, Protein Kinase, and Fungal Hydrophobin Activity

We detected several significantly enriched GO annotations across differentially expressed genes. Specifically, we found 26 enriched GO terms in the response to tolerance group and 66 enriched GO terms in the interaction effect. Conversely, we found no enriched GO terms in the response to Zn treatment. When considering genes differentially expressed between tolerant and sensitive isolates, there was a large overlap in the functions of enriched GO terms, so we merged them into four categories: transmembrane transport, oxidoreductase activity, fungal hydrophobins, and protein kinase activity. Transmembrane transport was the most enriched annotation category (highest % associated genes) and corresponded to three pleiotropic drug resistance (PDR) transporters (Figure 4). When considering genes differentially expressed in response to interaction effect, out of the four categories above, oxidoreductase activity was the only and most enriched annotation in this comparison (Supplemental Figure 1).

**Figure 4.**
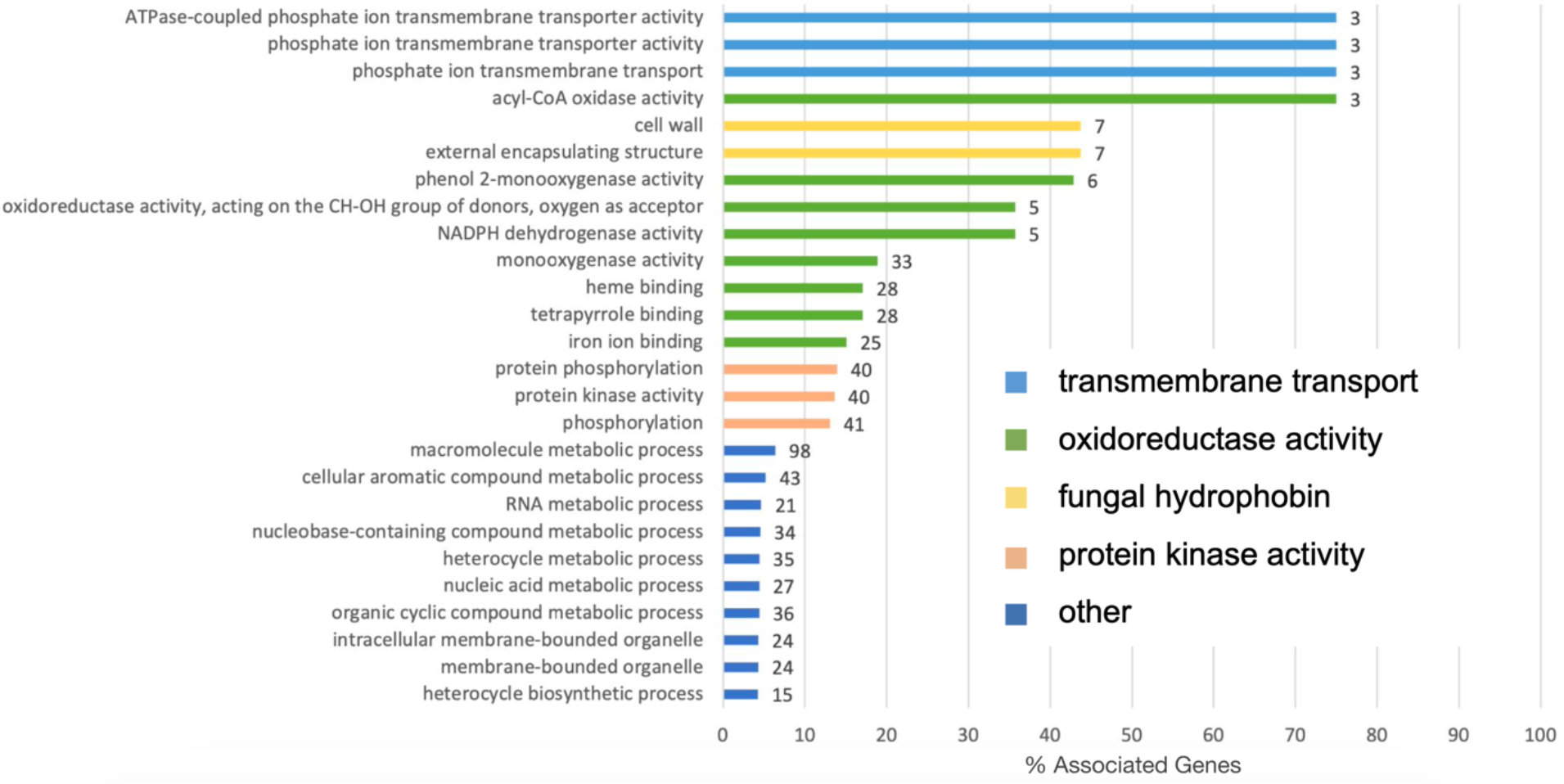
Significantly differentially expressed genes between tolerant and sensitive isolates are enriched in transmembrane transport, oxidoreductase, fungal hydrophobin, and protein kinase activity. Barplot showing enriched GO terms in significantly differentially expressed genes between tolerant and sensitive isolates. Bar length represents the percentage of all genes with the GO term annotation that appear in this gene group. The numbers at the end of bars represent the number of unique genes with that GO term annotation. Colors represent general functional categories (light blue = transmembrane transport, green = oxidoreductase activity, yellow = fungal hydrophobin, orange = protein kinase activity, dark blue = other).

### Differential Expression of Specific Genes of Interest

Most *S. luteus* previously identified Zn tolerance candidate genes (Zn transporters and genes genetically diverged between isolates from contaminated and noncontaminated sites, Ruytinx et al., 2017, Coninx et al., 2017, 2019, Bazzicalupo et al., 2020), showed significant differential expression (Table 1). Specifically, seven metal transport genes, two chelators, and two oxidative stress relief genes were differentially expressed between tolerant and sensitive isolates irrespective of Zn treatment (tolerance group FDR <0.05, interaction FDR > 0.05; Table 1A). Notably, none of the candidate genes showed significantly different gene expression across Zn treatment, irrespective of tolerance group (Zn treatment FDR <0.05, interaction FDR > 0.05; Table 1B). To highlight the interaction effect, we report the candidate genes differential expression results separately for tolerant and sensitive isolates (Table 1C-D). We found tolerant isolates to display no significant gene expression differences when exposed to Zn (Table 1B), suggesting these isolates were not stressed when exposed to 130 ppm Zn. Four Zn transporters, three metal chelation related genes, and two antioxidants were differently expressed between tolerant and sensitive isolates in response to Zn treatment. Specifically, two CDF transporters, one ZIP transporter, and three metal chelation related genes were upregulated in sensitive isolates in response to Zn and one CDF transporter and two antioxidants were downregulated in sensitive isolates in response to Zn (Table 1D).

**Table 1.**
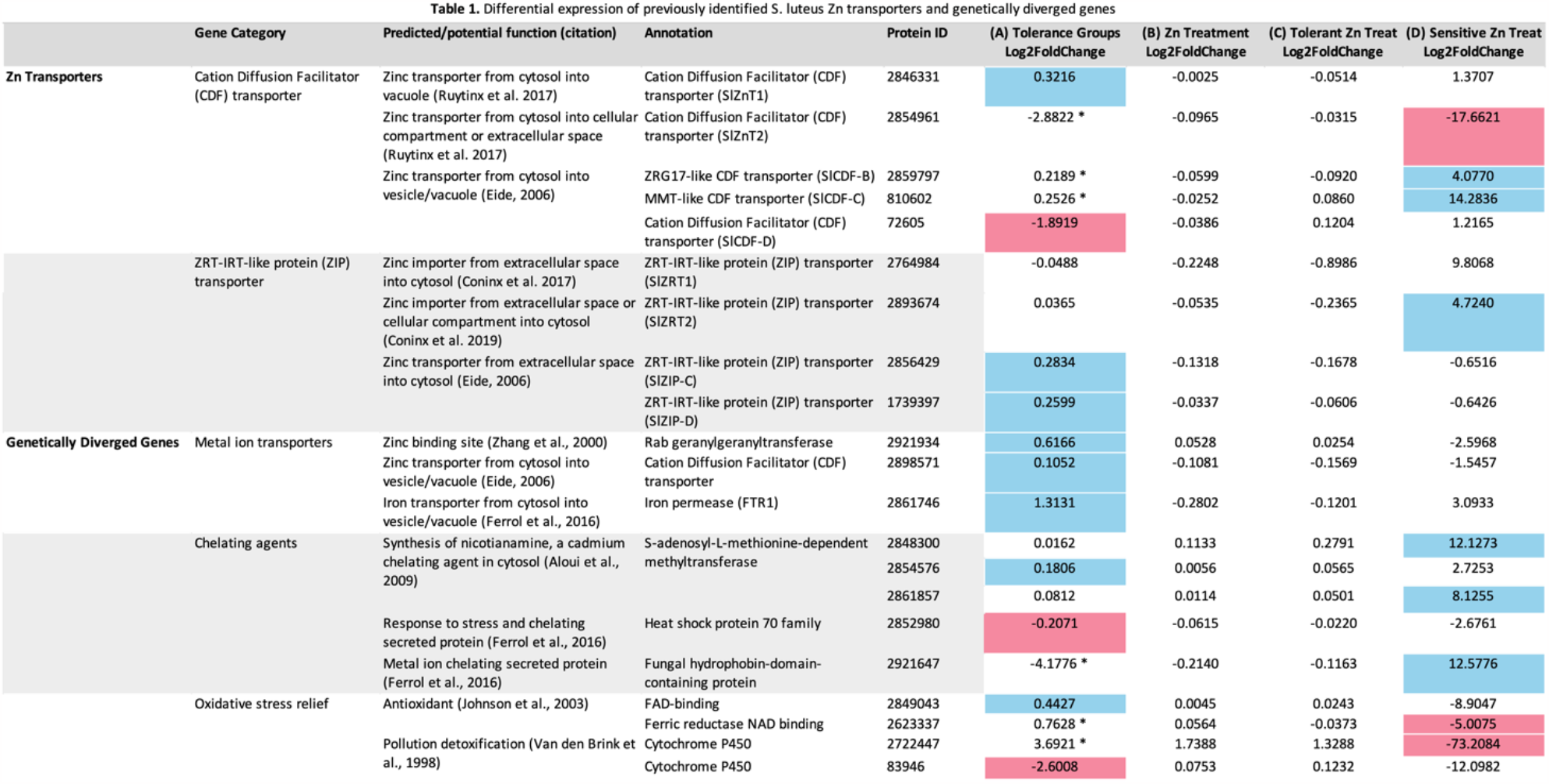
Differential expression of previously identified *S. luteus* Zn transporters and genetically diverged genes. Differential expression across model comparisons of previously identified genes putatively involved in *S. luteus* metal tolerance. Log fold change (lfc) values display differential gene expression in response tolerance groups **(A)**, to Zn treatment **(B)**, Zn treatment in tolerant isolates only **(C)**, and Zn treatment in sensitive isolates only **(D)**. Colored log fold change values have an adjusted p value of less than 0.05 and denote the direction of differential expression (blue = lfc > 0, red = lfc < 0). One star (*) depicts a significant log fold change value, but significant interaction effect supersedes main effects.

We found many other differentially expressed genes across each of the three model comparisons that have not been highlighted in previous studies. Of the genes with the most positive and negative log fold change values in the three model comparisons (n = 57), close to 70% had no annotation (Supplemental Table 3). The remaining annotated genes had predicted functions relating to processes such as nucleic acid metabolism (∼71% genes), signal transduction (∼6% genes), and fungal mating type (∼6% genes). Genes that were differentially expressed in response to Zn treatment had predicted functions related to signal transduction, oxidoreductase activity, transmembrane transport, and the regulation of nucleic acids (Supplemental Table 3).

## Discussion

We investigated Zn tolerance and differential gene expression in *S. luteus* isolates collected from metal contaminated and non-contaminated sites. We found that in this species Zn tolerance is associated with soil contamination, is both a constitutive and environmentally dependent trait, and results from a combination of responses involving metal exclusion and immobilization, as well as recognition and mitigation of metal-induced oxidative stress (summarized in Figure 5). Differentially expressed genes between tolerant and sensitive isolates were enriched in transmembrane transport, oxidoreductase, fungal hydrophobin, and protein kinase activity. We also found gene expression differences in previously studied candidate genes for *S. luteus* Zn tolerance. Even though Zn tolerant isolates showed no significant differences in candidate gene expression under high Zn, sensitive isolates displayed differential expression in candidate transporters, chelators, and antioxidants when exposed to Zn.

**Figure 5.**
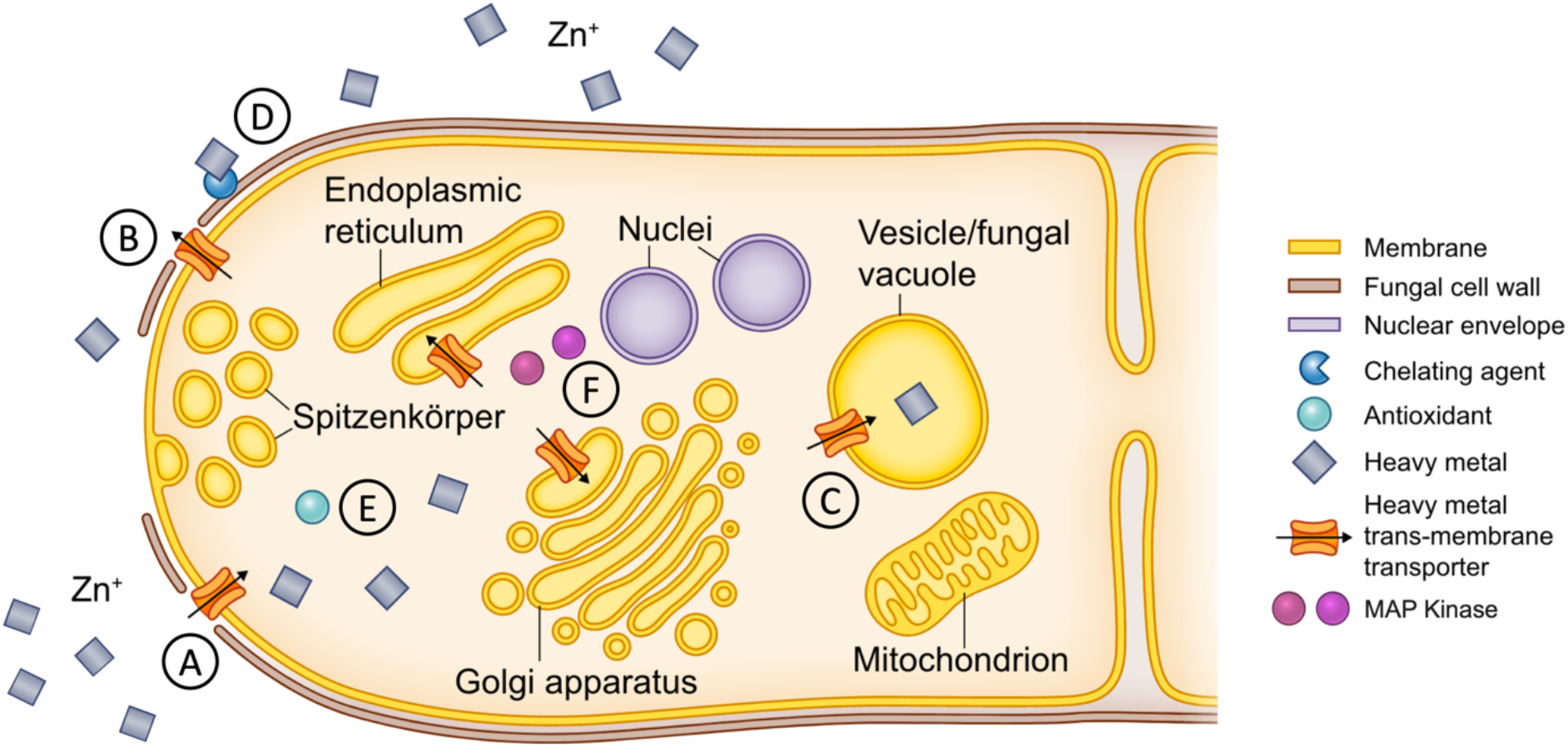
Potential mechanisms of Zn tolerance in *S. luteus*. Diagram of putative mechanisms of Zn tolerance in *S. luteus*. **(A-C)** Zn transmembrane transport including Zn importers (such as ZIP transporters), exporters (such as CDF transporters), and sequestration transporters (such as CDF and PDR genes) which maintain cytosolic Zn homeostasis. **(D)** Zinc chelation (for example through fungal hydrophobins) which immobilizes excess Zn. **(E)** Antioxidants which mitigate metal-induced oxidative stress. **(F)** MAP kinases which transmit stress signals to the nucleus. (Adapted from Branco et al., 2022)

We found *S. luteus* Zn tolerant isolates mainly associated with high soil Zn content, corroborating previous research (Colpaert et al, 2004). However, one isolate (LMD7) from the contaminated site showed Zn tolerance more similar to isolates from the noncontaminated site (Figure 1). This discrepancy is likely due to soil heterogeneity and the existence of low-Zn pockets within the contaminated site. In fact, the soil Zn dry weight concentrations at the contaminated site range from 1 to 197 ppm (Op De Beeck et al., 2015), making it likely for Zn sensitive fungi to be able to survive in localized areas of low Zn concentration. Another possibility is that soil Zn bioavailability varies across the contaminated site and that Zn can locally occur in a form that is not toxic to fungi, allowing one sensitive isolate to persist.

The *S. luteus* Zn tolerant and sensitive isolates showed markedly different overall transcriptomic profiles, with tolerant isolates displaying much higher overall transcriptomic variation (Figure 2). Notably, few individual genes showed differential expression in response to Zn treatment even though many genes were regulated differently in response to Zn between tolerant and sensitive isolates (interaction effect, Figure 3). A large number of differentially expressed genes showed a constitutive response (tolerance group effect), meaning that many genes potentially involved in *S. luteus* Zn tolerance are not differentially expressed in response to high Zn environments but instead Zn tolerant isolates consistently maintain a Zn tolerant gene expression profile. This pattern of constitutive expression independent of external Zn concentration had been previously found in two *S. luteus* Zn transmembrane transporters (Ruytinx et al., 2017) and we show here that it applies to a much larger number of genes. The large observed gene expression differences across tolerant isolates also suggests the existence of distinct paths to achieve Zn tolerance in the species, through the expression of different sets of metal tolerance related genes.

We found a clear enrichment of transmembrane transport, oxidoreductase, fungal hydrophobin, and protein kinase activity in response to tolerance group, corroborating marked differences across tolerant and sensitive isolates. These processes have been previously implicated in metal response (Wolfger et al., 2001, Bazzicalupo et al., 2020, Lin et al., 2005). Enriched transmembrane transport activity corresponded to three PDR transporters, ATP-binding cassette (ABC) transporters that have been linked to metal stress in both plants and fungi (Nuruzzaman et al., 2014, Wolfger et al., 2001). In yeast, two PDR transporters have been linked to vacuolar sequestration of chelated metal conjugates (Buechel and Pinkett, 2020) and could have a similar function in *S. luteus* (Figure 5). Oxidoreductase activity corresponded to around 50 genes in a variety of pathways that catalyze electron transfer between molecules (Bolann and Ulvik, 1997). Oxidative stress is a known byproduct of metal exposure and is characterized by the accumulation of reactive oxygen species (ROS) (Azevedo et al., 2007, Liang & Zhou., 2007). The buildup of ROS can lead to impaired cellular functioning and even cell death, so the reduction of ROS by oxidoreductase activity is an important homeostatic response (Zadrąg-Tęcza et al., 2018). Fungal hydrophobin activity corresponded to seven cell wall fungal hydrophobin genes that function by binding excess metals to render them inert (Ferrol et al., 2016, Figure 5) and have been implicated in the metal response of mycorrhizal fungi (Sammer et al., 2016). Protein kinase activity corresponded to close to 40 genes, many of which are predicted mitogen-activated protein kinases (MAPKs), evolutionary conserved signal transduction modules that convert environmental cues into cell responses (Mishra et al., 2006, Cristina et al., 2010). These proteins have been shown to be activated in response to metal stress (Lin et al., 2005, Mondal, 2022), to function in the signal cascade mechanism of metal homeostasis (Mondal, 2022), and likely play an important role in *S. luteus* Zn tolerance (Figure 5).

Differential expression of candidate genes across our three model comparisons further confirms marked differences across *S. luteus* Zn sensitive and tolerant isolates, affirms Zn tolerance in this species is comprised of constitutive differences and environmentally driven responses, and hints at potential *S. luteus* Zn tolerance mechanisms. Zn tolerance is partially achieved through constitutive differential expression in all mechanistic categories of the previously studied *S. luteus* candidate metal ion transporters, metal chelating agents, and antioxidants (Bazzicalupo et al., 2020, Table 1, Figure 5). In addition, Zn transporters, metal chelation related genes, and antioxidants were differently expressed between tolerant and sensitive isolates in response to Zn treatment (interaction effect). CDF transporters have been implied in the export and sequestration of excess Zn (Ruytinx et al., 2017) and their upregulation in high Zn environments is a potential mechanism of Zn tolerance in *S. luteus*. Chelating agents, such as fungal hydrophobins previously mentioned, act to bind and neutralize excess metals (Ferrol et al., 2016) and upregulation of these genes in high Zn environments is likely another mechanism of Zn tolerance in *S. luteus*. Conversely, the upregulation of a ZIP transporter (*SlZRT2*, 2893674), which functions in Zn import (Coninx et al., 2019), the downregulation of a CDF transporter (*SlZnT2*, 2854961), which functions in Zn export and sequestration (Ruytinx et al., 2017), and the downregulation of two antioxidants (2623337; 2722447), which alleviate oxidative stress caused by excess metal (Zadrąg-Tęcza et al., 2018), in sensitive isolates in high Zn are unexpected results. Potentially, the counterintuitive regulation of some of these genes could contribute to the Zn sensitivity of these isolates, though further study is required. Furthermore, Zn tolerant isolates showed no differences in candidate gene expression when exposed to Zn addition (Table 1). We suspect the amount of Zn added in our experiment was sufficiently high to induce a stress response in the sensitive but not the tolerant isolates. Future studies should consider including multiple levels of Zn treatment to ensure that tolerant isolates are being amply stressed. In summary, at the level of Zn treatment in our experiment, tolerant isolates seem to have been able to achieve Zn homeostasis with constitutive mechanisms alone, while sensitive isolates employed both constitutive mechanisms and environmental responses to survive in high Zn.

Even though the results reported here make significant contributions for understanding the mechanisms of fungal Zn tolerance, our study has some limitations that preclude unveiling additional processes potentially involved in tolerating high Zn. Specifically, it has previously been shown that ZIP transporters can have a time sensitive response to excess Zn, with downregulation lasting even only a few hours (Coninx et al. 2019). Since we extracted RNA after more than one week under Zn stress, the expression profile we captured here would have not reflected these immediate responses, but instead an adjusted equilibrium. Also, fungi most likely deal with excess levels of Zn in both acute and prolonged interactions so future work should examine the differences in the immediate responses of tolerant and sensitive *S. luteus* isolates to excess Zn. In addition, the available *S. luteus* reference genome annotation is far from complete, with well over 50% of genes missing a predicted function. This means that we likely missed important gene categories involved in Zn tolerance. Future studies should rely on an improved reference genome annotation and use higher metal concentrations to assess stress response in tolerant isolates.

In conclusion, our study unveils mechanisms of *S. luteus* Zn tolerance and contributes to the understanding of how fungi achieve metal tolerance. We found *S. luteus* displays constitutive differences and environmentally driven Zn responses. We also explored expression patterns of genes previously implicated in *S. luteus* metal tolerance and uncovered new signaling genes that potentially contribute to Zn tolerance in the species. Further research can address metal responses of mycorrhizal fungi in conjunction with their obligate plant partners, allowing further understanding of the mechanisms of metal tolerance in a more ecologically relevant context.

## Data Availability Statement

The sequence data generated and used in this study is available in the Sequence Read Archive (BioProject PRJNA972046-PRJNA972026). All analytical pipelines are available at https://github.com/ahsmith22/SluteusRNA.

## Funding Statement

This work was supported by JGI CPSNI 506158 awarded to SB, H-LL, and JR, NSF IOS-PBI (2029168) awarded to SB and H-LL, Research Foundation Flanders FWO G011723N and VUB OZR3483 awarded to JR. The work (proposal: 10.46936/10.25585/60000469) conducted by the U.S. Department of Energy Joint Genome Institute (https://ror.org/04xm1d337), a DOE Office of Science User Facility, is supported by the Office of Science of the U.S. Department of Energy operated under Contract No. DE-AC02-05CH11231.

## Acknowlegements

We thank Jessica Fletcher for comments on earlier drafts.

## Notes

https://github.com/ahsmith22/SluteusRNA

### Competing Interest Statement

The authors have declared no competing interest.

## References

Anahid, S., Yaghmaei, S., Ghobadinejad, Z., 2011. Heavy metal tolerance of fungi. Sci. Iran. 18, 502–508. 10.1016/j.scient.2011.05.015

Azevedo, M.-M., Carvalho, A., Pascoal, C., Rodrigues, F., Cássio, F., 2007. Responses of antioxidant defenses to Cu and Zn stress in two aquatic fungi. Sci. Total Environ. 377, 233–243. 10.1016/j.scitotenv.2007.02.027

Bazzicalupo, A.L., Ruytinx, J., Ke, Y.-H., Coninx, L., Colpaert, J.V., Nguyen, N.H., Vilgalys, R., Branco, S., 2020. Fungal heavy metal adaptation through single nucleotide polymorphisms and copy-number variation. Mol. Ecol. 29, 4157–4169. 10.1111/mec.15618

Bellion, M., Courbot, M., Jacob, C., Blaudez, D., Chalot, M., 2006. Extracellular and cellular mechanisms sustaining metal tolerance in ectomycorrhizal fungi. FEMS Microbiol. Lett. 254, 173–181. 10.1111/j.1574-6968.2005.00044.x

Bindea, G., Mlecnik, B., Hackl, H., Charoentong, P., Tosolini, M., Kirilovsky, A., Fridman, W.-H., Pagès, F., Trajanoski, Z., Galon, J., 2009. ClueGO: a Cytoscape plug-in to decipher functionally grouped gene ontology and pathway annotation networks. Bioinformatics 25, 1091–1093. 10.1093/bioinformatics/btp101

Blaudez, D., Chalot, M., 2011. Characterization of the ER-located zinc transporter ZnT1 and identification of a vesicular zinc storage compartment in Hebeloma cylindrosporum. Fungal Genet. Biol. 48, 496–503. 10.1016/j.fgb.2010.11.007

Bolann, B.J., Ulvik, R.J., 1997. How do antioxidants work? Tidsskrift for den Norske Lægeforening 117, 1928–1932. PMID: 9214016.

Branco, S., Schauster, A., Liao, H.-L., Ruytinx, J., 2022. Mechanisms of stress tolerance and their effects on the ecology and evolution of mycorrhizal fungi. New Phytol 235, 2158–2175. 10.1111/nph.18308

Buechel, E.R., Pinkett, H.W., 2020. Transcription factors and ABC transporters: from pleiotropic drug resistance to cellular signaling in yeast. FEBS Letters 594, 3943–3964. 10.1002/1873-3468.13964

Bushnell, B., 2014. BBMap: BBMap short read aligner, and other bioinformatic tools. https://sourceforge.net/projects/bbmap/

Colpaert, J.V., 2008. Chapter 11 Heavy metal pollution and genetic adaptations in ectomycorrhizal fungi. British Mycological Society Symposia Series, Stress in Yeast and Filamentous Fungi 27, 157–173. 10.1016/S0275-0287(08)80053-7

Colpaert, J.V., Adriaensen, K., Muller, L.A.H., Lambaerts, M., Faes, C., Carleer, R., Vangronsveld, J., 2005. Element profiles and growth in Zn-sensitive and Zn-resistant Suilloid fungi. Mycorrhiza 15, 628–634. 10.1007/s00572-005-0009-6

Colpaert, J.V., Muller, L.A.H., Lambaerts, M., Adriaensen, K., Vangronsveld, J., 2004. Evolutionary Adaptation to Zn Toxicity in Populations of Suilloid Fungi. New Phytol 162, 549–559. 10.1111/j.1469-8137.2004.01037.x

Colpaert, J.V., van Assche, J.A., 1987. Heavy Metal Tolerance in Some Ectomycorrhizal Fungi. Funct. Ecol. 1, 415–421. 10.2307/2389799

Colpaert, J.V., Vandenkoornhuyse, P., Adriaensen, K., Vangronsveld, J., 2000. Genetic Variation and Heavy Metal Tolerance in the Ectomycorrhizal Basidiomycete Suillus luteus. New Phytol 147, 367–379. 10.1046/j.1469-8137.2000.00694.x

Coninx, L., Smisdom, N., Kohler, A., Arnauts, N., Ameloot, M., Rineau, F., Colpaert, J.V., Ruytinx, J., 2019. SlZRT2 Encodes a ZIP Family Zn Transporter With Dual Localization in the Ectomycorrhizal Fungus Suillus luteus. Front. Microbiol. 10, 2251. 10.3389/fmicb.2019.02251

Coninx, L., Thoonen, A., Slenders, E., Morin, E., Arnauts, N., Op De Beeck, M., Kohler, A., Ruytinx, J., Colpaert, J.V., 2017. The SlZRT1 Gene Encodes a Plasma Membrane-Located ZIP (Zrt-, Irt-Like Protein) Transporter in the Ectomycorrhizal Fungus Suillus luteus. Front. Microbiol. 8, 2320. 10.3389/fmicb.2017.02320

Cristina, M., Petersen, M., Mundy, J., 2010. Mitogen-Activated Protein Kinase Signaling in Plants. Annu. Rev. Plant Biol. 61, 621–649. 10.1146/annurev-arplant-042809-112252

Ezzouhri, L., Castro, E., Moya, M., Espinola, F., Lairini, K., 2009. Heavy metal tolerance of filamentous fungi isolated from polluted sites in Tangier, Morocco. Afr. J. microbiol. Res. 3. 10.5897/AJMR.9000354

Faggioli, V., Menoyo, E., Geml, J., Kemppainen, M., Pardo, A., Salazar, M.J., Becerra, A.G., 2019. Soil lead pollution modifies the structure of arbuscular mycorrhizal fungal communities. Mycorrhiza 29, 363–373. 10.1007/s00572-019-00895-1

Feldmann, H., 2012. Yeast: molecular and cell biology, 2nd complete rev. and greatly enl. ed. ed. John Wiley, Hoboken.

Ferrol, N., Tamayo, E., Vargas, P., 2016. The heavy metal paradox in arbuscular mycorrhizas: from mechanisms to biotechnological applications. JXB 67, 6253–6265. 10.1093/jxb/erw403

Galván Márquez, I., Ghiyasvand, M., Massarsky, A., Babu, M., Samanfar, B., Omidi, K., Moon, T.W., Smith, M.L., Golshani, A., 2018. Zinc oxide and silver nanoparticles toxicity in the baker’s yeast, Saccharomyces cerevisiae. PLoS ONE 13, e0193111. 10.1371/journal.pone.0193111

Guo, K., Liu, Y. f., Zeng, C., Chen, Y. y., Wei, X. j., 2014. Global research on soil contamination from 1999 to 2012: A bibliometric analysis. Acta Agric Scand B Soil Plant Sci. 64, 377–391. 10.1080/09064710.2014.913679

Howe, R., Evans, R.L., Ketteridge, S.W., 1997. Copper-binding proteins in ectomycorrhizal fungi. New Phytol 135, 123–131. 10.1046/j.1469-8137.1997.00622.x

Khouja, H.R., Abbà, S., Lacercat-Didier, L., Daghino, S., Doillon, D., Richaud, P., Martino, E., Vallino, M., Perotto, S., Chalot, M., Blaudez, D., 2013. OmZnT1 and OmFET, two metal transporters from the metal-tolerant strain Zn of the ericoid mycorrhizal fungus Oidiodendron maius, confer zinc tolerance in yeast. Fungal Genet. Biol. 52, 53–64. 10.1016/j.fgb.2012.11.004

Kim, D., Langmead, B., Salzberg, S.L., 2015. HISAT: a fast spliced aligner with low memory requirements. Nat Methods 12, 357–360. 10.1038/nmeth.3317

Kohler, A., Kuo, A., Nagy, L.G., Morin, E., Barry, K.W., Buscot, F., Canbäck, B., Choi, C., Cichocki, N., Clum, A., Colpaert, J., Copeland, A., Costa, M.D., Doré, J., Floudas, D., Gay, G., Girlanda, M., Henrissat, B., Herrmann, S., Hess, J., Högberg, N., Johansson, T., Khouja, H.-R., LaButti, K., Lahrmann, U., Levasseur, A., Lindquist, E.A., Lipzen, A., Marmeisse, R., Martino, E., Murat, C., Ngan, C.Y., Nehls, U., Plett, J.M., Pringle, A., Ohm, R.A., Perotto, S., Peter, M., Riley, R., Rineau, F., Ruytinx, J., Salamov, A., Shah, F., Sun, H., Tarkka, M., Tritt, A., Veneault-Fourrey, C., Zuccaro, A., Tunlid, A., Grigoriev, I.V., Hibbett, D.S., Martin, F., 2015. Convergent losses of decay mechanisms and rapid turnover of symbiosis genes in mycorrhizal mutualists. Nat Genet 47, 410–415. 10.1038/ng.3223

Lanfranco, L., Balsamo, R., Martino, E., Perotto, S., Bonfante, P., 2010. Zinc ions alter morphology and chitin deposition in an ericoid fungus. Eur J Histochem 46, 341. 10.4081/1746

Li, C., Zhou, K., Qin, W., Tian, C., Qi, M., Yan, X., Han, W., 2019. A Review on Heavy Metals Contamination in Soil: Effects, Sources, and Remediation Techniques. Soil Sediment Contam 28, 380–394. 10.1080/15320383.2019.1592108

Liang, Q., Zhou, B., 2007. Copper and Manganese Induce Yeast Apoptosis via Different Pathways. MBoC 18, 4741–4749. 10.1091/mbc.e07-05-0431

Liao, Y., Smyth, G.K., Shi, W., 2014. featureCounts: an efficient general purpose program for assigning sequence reads to genomic features. Bioinformatics 30, 923–930. 10.1093/bioinformatics/btt656

Lin, C.-W., Chang, H.-B., Huang, H.-J., 2005. Zinc induces mitogen-activated protein kinase activation mediated by reactive oxygen species in rice roots. Plant Physiol. Biochem. 43, 963–968. 10.1016/j.plaphy.2005.10.001

Lofgren, L., Nguyen, N., Kennedy, P., Perez-Pazos, E., Fletcher, J., Liao, H.-L., Wang, H., Zhang, K., Ruytinx, J., Smith, A. H., Yi-Hong, K., Cotter, H. Van T., Engwall, E. Hameed, K., Vilgalys, R., Branco, S. In review. New Phytologist.

Love, M.I., Huber, W., Anders, S., 2014. Moderated estimation of fold change and dispersion for RNA-seq data with DESeq2. Genome Biol. 15, 550. 10.1186/s13059-014-0550-8

Martino, E., Perotto, S., Parsons, R., Gadd, G.M., 2003. Solubilization of insoluble inorganic zinc compounds by ericoid mycorrhizal fungi derived from heavy metal polluted sites. Soil Biol. Biochem. 35, 133–141. 10.1016/S0038-0717(02)00247-X

Miransari, M., 2011. Hyperaccumulators, arbuscular mycorrhizal fungi and stress of heavy metals. Biotechnol. Adv. 29, 645–653. 10.1016/j.biotechadv.2011.04.006

Mishra, N.S., Tuteja, R., Tuteja, N., 2006. Signaling through MAP kinase networks in plants. Arch. Biochem. Biophys. 452, 55–68. 10.1016/j.abb.2006.05.001

Mondal, S., 2022. Heavy Metal Stress–Induced Activation of Mitogen-Activated Protein Kinase Signalling Cascade in Plants. Plant Mol Biol Rep. 10.1007/s11105-022-01350-w

Nuruzzaman, M., Zhang, R., Cao, H.-Z., Luo, Z.-Y., 2014. Plant Pleiotropic Drug Resistance Transporters: Transport Mechanism, Gene Expression, and Function. J. Integr. Plant Biol. 56, 729–740. 10.1111/jipb.12196

Oksanen, J., Simpson, G., Blanchet, F., Kindt, R., Legendre, P., Minchin, P., O’Hara, R., Solymos, P., Stevens, M., Szoecs, E., Wagner, H., Barbour, M., Bedward, M., Bolker, B., Borcard, D., Carvalho, G., Chirico, M., De Caceres, M., Durand, S., Evangelista, H., FitzJohn, R., Friendly, M., Furneaux, B., Hannigan, G., Hill, M., Lahti, L., McGlinn, D., Ouellette, M., Ribeiro Cunha, E., Smith, T., Stier, A., Ter Braak, C., Weedon, J., 2022. vegan: Community Ecology Package. R package Version 2.4-6. https://CRAN.R-project.org/package=vegan

Op De Beeck, M., Ruytinx, J., Smits, M.M., Vangronsveld, J., Colpaert, J.V., Rineau, F., 2015. Belowground fungal communities in pioneer Scots pine stands growing on heavy metal polluted and non-polluted soils. Soil Biol. Biochem. 86, 58–66. 10.1016/j.soilbio.2015.03.007

Påhlsson, A.-M.B., 1989. Toxicity of heavy metals (Zn, Cu, Cd, Pb) to vascular plants. Water Air Soil Pollut 47, 287–319. https://doi-org.aurarialibrary.idm.oclc.org/10.1007/BF00279329

Pawlowska, T.E., Charvat, I., 2004. Heavy-Metal Stress and Developmental Patterns of Arbuscular Mycorrhizal Fungi. Appl Environ Microbiol 70, 6643–6649. 10.1128/AEM.70.11.6643-6649.2004

Priyadarshini, E., Priyadarshini, S.S., Cousins, B.G., Pradhan, N., 2021. Metal-Fungus interaction: Review on cellular processes underlying heavy metal detoxification and synthesis of metal nanoparticles. Chemosphere 274, 129976. 10.1016/j.chemosphere.2021.129976

R Core Team, 2020. R: A language and environment for statistical computing. https://www.R-project.org/

Ramrakhiani, L., Halder, A., Majumder, A., Mandal, A.K., Majumdar, S., Ghosh, S., 2017. Industrial waste derived biosorbent for toxic metal remediation: Mechanism studies and spent biosorbent management. J. Chem. Eng. 308, 1048–1064. 10.1016/j.cej.2016.09.145

Ritz, C., Baty, F., Streibig, J.C., Gerhard, D., 2015. Dose-Response Analysis Using R. PLOS ONE 10, e0146021. 10.1371/journal.pone.0146021

Robinson, J.R., Isikhuemhen, O.S., Anike, F.N., 2021. Fungal–Metal Interactions: A Review of Toxicity and Homeostasis. J. Fungus 7, 225. 10.3390/jof7030225

RStudio Team, 2019. RStudio: Integrated Development for R. http://www.rstudio.com/

Ruytinx, J., Coninx, L., Nguyen, H., Smisdom, N., Morin, E., Kohler, A., Cuypers, A., Colpaert, J.V., 2017. Identification, evolution and functional characterization of two Zn CDF-family transporters of the ectomycorrhizal fungus Suillus luteus. Environ. Microbiol. Rep. 9, 419–427. 10.1111/1758-2229.12551

Ruytinx, J., Craciun, A.R., Verstraelen, K., Vangronsveld, J., Colpaert, J.V., Verbruggen, N., 2011. Transcriptome analysis by cDNA-AFLP of Suillus luteus Cd-tolerant and Cd-sensitive isolates. Mycorrhiza 21, 145–54. 10.1007/s00572-010-0318-2

Sammer, D., Krause, K., Gube, M., Wagner, K., Kothe, E., 2016. Hydrophobins in the Life Cycle of the Ectomycorrhizal Basidiomycete Tricholoma vaccinum. PLOS ONE 11, e0167773. 10.1371/journal.pone.0167773

Shannon, P., Markiel, A., Ozier, O., Baliga, N.S., Wang, J.T., Ramage, D., Amin, N., Schwikowski, B., Ideker, T., 2003. Cytoscape: a software environment for integrated models of biomolecular interaction networks. Genome Res 13, 2498–2504. 10.1101/gr.1239303

Sharma, A., Patni, B., Shankhdhar, D., Shankhdhar, S.C., 2013. Zinc – An Indispensable Micronutrient. Physiol Mol Biol Plants 19, 11–20. 10.1007/s12298-012-0139-1

Smith, P., House, J.I., Bustamante, M., Sobocká, J., Harper, R., Pan, G., West, P.C., Clark, J.M., Adhya, T., Rumpel, C., Paustian, K., Kuikman, P., Cotrufo, M.F., Elliott, J.A., McDowell, R., Griffiths, R.I., Asakawa, S., Bondeau, A., Jain, A.K., Meersmans, J., Pugh, T.A.M., 2016. Global change pressures on soils from land use and management. Glob Change Biol 22, 1008–1028. 10.1111/gcb.13068

Sonke, J.E., Hoogewerff, J.A., van der Laan, S.R., Vangronsveld, J., 2002. A chemical and mineralogical reconstruction of Zn-smelter emissions in the Kempen region (Belgium), based on organic pool sediment cores. Sci. Total Environ. 292, 101–119. 10.1016/S0048-9697(02)00033-5

Teng, Y., Du, X., Wang, T., Mi, C., Yu, H., Zou, L., 2018. Isolation of a fungus Pencicillium sp. with zinc tolerance and its mechanism of resistance. Arch Microbiol 200, 159–169. 10.1007/s00203-017-1430-x

The UniProt Consortium, 2023. UniProt: the Universal Protein Knowledgebase in 2023. Nucleic Acids Res 51, D523–D531. 10.1093/nar/gkac1052

Wang, A.S., Angle, J.S., Chaney, R.L., Delorme, T.A., Reeves, R.D., 2006. Soil pH Effects on Uptake of Cd and Zn by Thlaspi caerulescens. Plant Soil 281, 325–337. 10.1007/s11104-005-4642-9

Wolfger, H., Mamnun, Y.M., Kuchler, K., 2001. Fungal ABC proteins: pleiotropic drug resistance, stress response and cellular detoxification. Res. Microbiol. 152, 375–389. 10.1016/S0923-2508(01)01209-8

Zadrąg-Tęcza, R., Maślanka, R., Bednarska, S., Kwolek-Mirek, M., 2018. Response Mechanisms to Oxidative Stress in Yeast and Filamentous Fungi, in: Skoneczny, M. (Ed.), Stress Response Mechanisms in Fungi: Theoretical and Practical Aspects. Springer International Publishing, Cham, pp. 1–34. 10.1007/978-3-030-00683-9_1

