## Supplemental Materials for "Comparative transcriptomics provides insight into molecular mechanisms of zinc tolerance in the ectomycorrhizal fungus *Suillus luteus*"

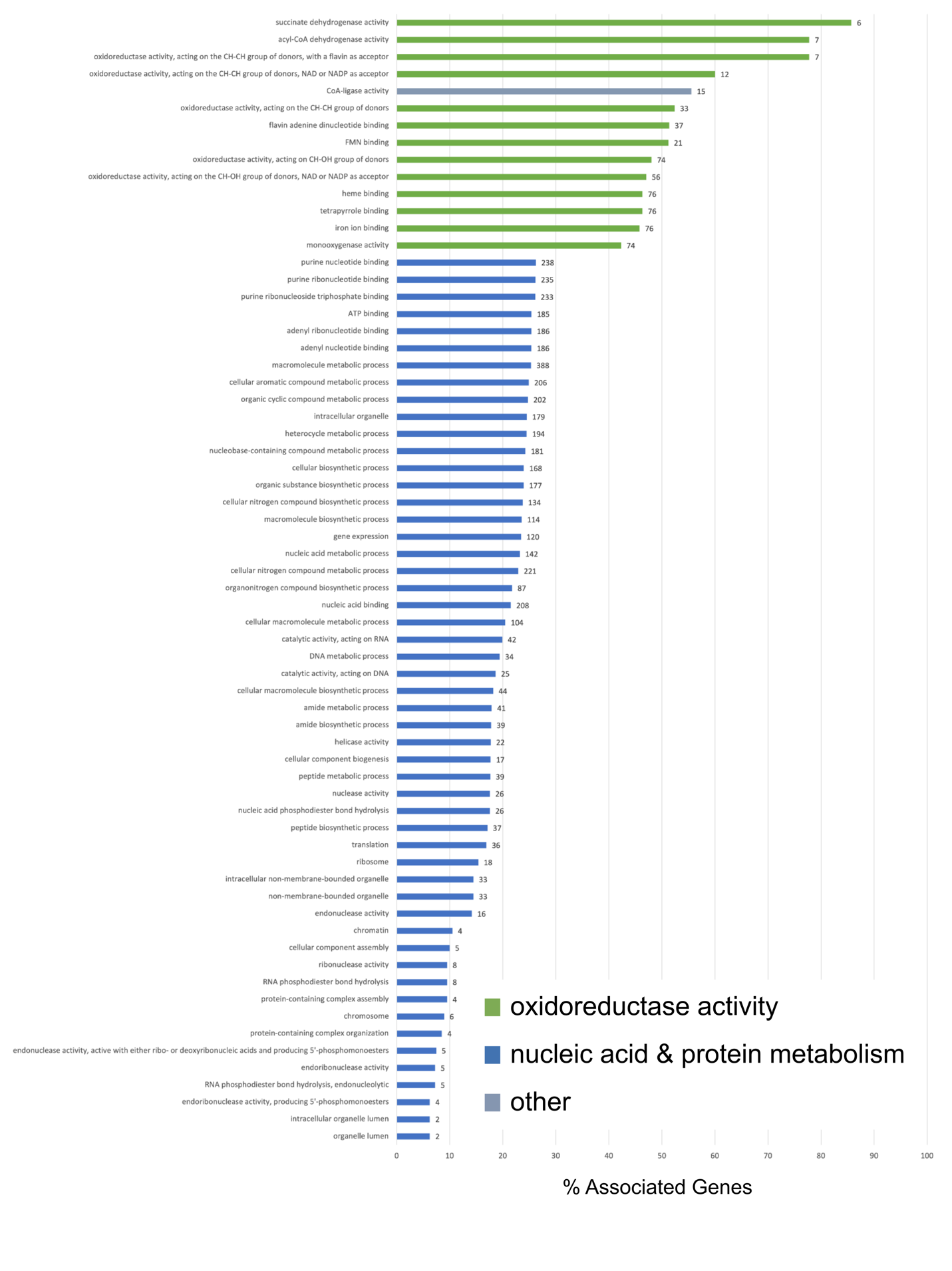

**Supplemental Figure 1. Significantly differentially expressed genes in the interaction effect are enriched in protein kinase and oxidoreductase activity**

Barplot showing enriched GO terms in significantly differentially expressed genes in the interaction effect. Bar length represents the percentage of all genes with the GO term annotation that appear in this gene group. The numbers at the end of bars represent the number of unique genes with that GO term annotation. Colors represent general functional categories (green = oxidoreductase activity, orange = protein kinase activity, blue = other).

**Supplemental Table 1.** *S. luteus* isolates included in the experiment, including collection year, Zn tolerance, and location of origin.

| Sample Name | Collection year | EC50 Zn (ppm) | Location |
| --- | --- | --- | --- |
| UH-Slu-Lm3 | 2000 | 396.10 | Lommel, BE |
| UH-Slu-Lm5 | 2000 | 641.84 | Lommel, BE |
| UH-Slu-Lm12 | 2000 | 936.98 | Lommel, BE |
| UH-Slu-LmD7 | 2011 | 142.78 | Lommel, BE |
| UH-Slu-Sl20 | 1998 | 573.91 | Lommel, BE |
| UH-Slu-P1 | 2000 | 61.34 | Paal, BE |
| UH-Slu-P2 | 2000 | 74.97 | Paal, BE |
| UH-Slu-P13 | 2000 | 77.06 | Paal, BE |
| UH-Slu-PD13 | 2011 | 207.78 | Paal, BE |
| UH-Slu-PD14 | 2011 | 103.93 | Paal, BE |

| **Supplemental Table 2.** Fries Media Recipe | |
| --- | --- |
| **Compound** | **Concentration** |
| ammonium tartrate | 5.4 mM |
| KH_2_PO_4_ | 1.5 mM |
| MgSO_4_·7H_2_O | 0.4 mM |
| NaCl | 0.3 mM |
| CaCl_2_·2H_2_O | 0.2 mM |
| FeCl_3_·6H_2_O | 4 μM |
| thiamine-HCl | 0.3 μM |
| MnSO_4_·H_2_O | 6 μM |
| CuSO_4_·5H_2_O | 0.8 μM |
| myo-inositol | 56 μM |
| biotin | 0.1 μM |
| pyridoxine | 0.5 μM |
| riboflavin | 0.3 μM |
| nicotinamide | 0.8 μM |
| p-aminobenzoic acid | 0.7 μM |
| Ca-pantothenate | 0.2 μM |
| glucose | 28 mM |
| agar | 0.8% |

**Supplemental Table 3.** Genes of Interest

| **Most Positively and Negatively Differentially Expressed Genes** | | | | |
| --- | --- | --- | --- | --- |
| **contrast** | **proteinID** | **lfc** | **GOterms** | **other annotations** |
| Inte | 2870763 | -242.4274 |  | Integrase/recombinase, N-terminal |
| Inte | 1883923 | -235.4473 | GUF |  |
| Inte | 2838289 | -230.6258 | GUF |  |
| Inte | 2054458 | -226.6317 | DNA binding | Integrase/recombinase, N-terminal |
| Inte | 2365220 | -223.4095 | GUF |  |
| Inte | 2761021 | -218.5840 | DNA binding | DNA breaking-rejoining enzyme, catalytic core |
| Inte | 2915549 | -214.1998 | GUF |  |
| Inte | 1762400 | -210.0848 | DNA binding | DNA breaking-rejoining enzyme, catalytic core |
| Inte | 2841028 | -206.2956 | GUF |  |
| Inte | 2601728 | -200.9169 | GUF |  |
| Inte | 2403967 | 217.9362 | GUF |  |
| Inte | 2925302 | 211.7357 | GUF |  |
| Inte | 1838360 | 209.2458 | GUF |  |
| Inte | 99884 | 205.3912 |  | Integrase/recombinase, N-terminal |
| Inte | 2911658 | 205.3313 | GUF |  |
| Inte | 2913934 | 196.0828 | GUF |  |
| Inte | 2837180 | 194.3027 | GUF |  |
| Inte | 2925322 | 189.9408 | GUF |  |
| Inte | 2955693 | 188.8173 | GUF |  |
| Inte | 16304 | 187.0048 | GUF |  |
| TvSe | 2854629 | -30.3685 | GUF |  |
| TvSe | 245087 | -30.3403 | GUF |  |
| TvSe | 2921847 | -30.3211 | GUF |  |
| TvSe | 2860109 | -29.2589 |  | Chromodomain-helicase DNA-binding protein |
| TvSe | 2772564 | -28.5703 | RNA-DNA hybrid ribonuclease activity |  |
| TvSe | 2923432 | -27.3839 | ATP binding | ATPase |
| TvSe | 2220471 | -26.6458 | GUF |  |
| TvSe | 2812508 | -25.2019 | GUF |  |
| TvSe | 2856202 | -24.9691 | GUF |  |
| TvSe | 32641 | -24.2737 | protein kinase activity | Protein tyrosine and serine/threonine kinase |
| TvSe | 2834385 | 21.7037 | GUF |  |
| TvSe | 1435914 | 21.1006 | GUF |  |
| TvSe | 567254 | 20.5787 | DNA binding | helix-turn-helix, Psq domain |
| TvSe | 2894618 | 16.1038 | GUF |  |
| TvSe | 2895427 | 14.7871 | GUF |  |
| TvSe | 2872549 | 14.7297 | GUF |  |
| TvSe | 2859448 | 14.0525 | GUF |  |
| TvSe | 808753 | 13.6615 | GTP binding | 50S ribosome-binding GTPase |
| Zne | 1863499 | -13.3096 | GUF |  |
| Zne | 2917334 | -12.3377 | GUF |  |
| Zne | 93300 | -12.3186 | metal ion binding | FOG: Zn-finger |
| Zne | 1754956 | -12.2716 | GUF |  |
| Zne | 2966410 | -12.2052 |  | N-methyltransferase |
| Zne | 1838502 | -12.1751 | GUF |  |
| Zne | 1753783 | -12.0731 | GUF |  |
| Zne | 1886202 | -12.0516 | zinc ion binding | Predicted E3 ubiquitin ligase |
| Zne | 15038 | -12.0331 | GUF |  |
| Zne | 2743056 | -12.0174 | GUF |  |
| Zne | 477002 | 6.7908 | GUF |  |
| Zne | 2107445 | 6.3020 | GUF |  |
| Zne | 814600 | 5.0389 | GUF |  |
| Zne | 12475 | 4.6136 | GUF |  |
| Zne | 2354186 | 3.0622 | protein binding | Heterokaryon incompatibility protein (HET) |
| Zne | 2783347 | 2.6967 | tryptophan biosynthetic process | Anthranilate phosphoribosyltransferase |
| Zne | 2849780 | 2.6669 | protein binding | F-box-like |
| Zne | 2918252 | 2.5984 | GUF |  |
| Zne | 1488136 | 2.5620 | GUF |  |
| **Genes Significantly Differentially Expressed in Response to Zn Treatment** | | | | |
|  | **proteinID** | **lfc** | **GOterms** | **other annotations** |
|  | 2861537 | -11.5660 | GUF |  |
|  | 108662 | -11.3553 |  | SAP Domain |
|  | 90983 | -10.8890 | protein binding | WD domain, G-beta repeat |
|  | 2916564 | -9.0037 | GUF |  |
|  | 2925319 | -8.8966 | GUF |  |
|  | 267929 | -7.1964 | GUF |  |
|  | 2837919 | -6.7521 |  | Protein kinase |
|  | 2852888 | -5.0391 | protein kinase activity | dual-specificity kinase |
|  | 2961255 | -3.4578 | DNA binding | Winged helix-turn helix |
|  | 2764313 | -3.4122 | DNA binding | Integrase/recombinase, N-terminal |
|  | 2338270 | -3.1998 | GUF |  |
|  | 810429 | -3.0438 | C-terminal protein methylation | Isoprenylcysteine carboxyl methyltransferase (ICMT) family |
|  | 2537479 | -2.9848 | GUF |  |
|  | 2873673 | -2.9663 | GUF |  |
|  | 2338282 | -2.5971 | GUF |  |
|  | 102388 | -2.3226 |  | P-loop containing nucleoside triphosphate hydrolase |
|  | 2852815 | -2.2723 | GUF |  |
|  | 2408646 | -2.0099 | protein binding | BTB/POZ domain |
|  | 2978110 | -1.9198 |  | P-loop containing nucleoside triphosphate hydrolase |
|  | 810904 | -1.5664 | GUF |  |
|  | 568711 | -0.6838 | GUF |  |
|  | 2865753 | 0.4686 | hydrolase activity; metal ion binding | Arginase family protein |
|  | 2977515 | 0.6452 | protein binding | Glutathione S-transferase, N-terminal domain |
|  | 2865726 | 0.6719 | GUF |  |
|  | 2862693 | 1.2714 | transmembrane transport | Sugar (and other) transporter |
|  | 809705 | 1.3017 | GUF |  |
|  | 156297 | 1.6658 | GUF |  |
|  | 2847486 | 1.6859 | GUF |  |
|  | 19649 | 1.7142 | GUF |  |
|  | 2840485 | 1.8286 | GUF |  |
|  | 2753246 | 1.8883 | GUF |  |
|  | 2868654 | 1.9739 | NAD+ kinase activity | ATP-NAD kinase N-terminal domain |
|  | 804592 | 2.0337 | GUF |  |
|  | 2841648 | 2.0949 | GUF |  |
|  | 812496 | 2.1300 | GUF |  |
|  | 107730 | 2.1725 | GUF |  |
|  | 2985300 | 2.2785 | nucleobase transmembrane transporter activity | Uridine permease/thiamine transporter/allantoin transport |
|  | 2919544 | 2.2920 | protein binding | F-box-like |
|  | 2923243 | 2.3786 | GUF |  |

*GUF = Gene of Unknown Function

**Supplemental Table 4.** SRA Database codes for all samples

| Sequencing Project ID | Library Name | BioProject | BioSample | Project Accession | Run Accession | Load Date |
| --- | --- | --- | --- | --- | --- | --- |
| 1354374 | HZGYU | PRJNA972046 | SAMN35066492 | SRP442734 | SRR24892960 | 2023-06-11 16:39:04 |
| 1354374 | HZGYW | PRJNA972047 | SAMN35066465 | SRP442733 | SRR24892961 | 2023-06-11 16:39:04 |
| 1354374 | HZGYX | PRJNA972048 | SAMN35066455 | SRP442732 | SRR24892962 | 2023-06-11 16:39:04 |
| 1354374 | HZGYY | PRJNA972049 | SAMN35066402 | SRP442731 | SRR24892963 | 2023-06-11 16:39:04 |
| 1354374 | HZGYZ | PRJNA972050 | SAMN35066497 | SRP442730 | SRR24892964 | 2023-06-11 16:39:04 |
| 1354374 | HZGZA | PRJNA972051 | SAMN35066498 | SRP442729 | SRR24892965 | 2023-06-11 16:39:04 |
| 1354375 | HZGZB | PRJNA972052 | SAMN35066464 | SRP442758 | SRR24893006 | 2023-06-11 16:39:41 |
| 1354375 | HZGZC | PRJNA972053 | SAMN35066445 | SRP442761 | SRR24893005 | 2023-06-11 16:39:41 |
| 1354375 | HZGZG | PRJNA972054 | SAMN35066456 | SRP442757 | SRR24893007 | 2023-06-11 16:39:41 |
| 1354375 | HZGZH | PRJNA972055 | SAMN35066403 | SRP442756 | SRR24893008 | 2023-06-11 16:39:41 |
| 1354375 | HZGZN | PRJNA972056 | SAMN35066446 | SRP442754 | SRR24893010 | 2023-06-11 16:39:41 |
| 1354375 | HZGZO | PRJNA972057 | SAMN35066482 | SRP442755 | SRR24893009 | 2023-06-11 16:39:41 |
| 1354376 | HZGZP | PRJNA972058 | SAMN35066460 | SRP442781 | SRR24893027 | 2023-06-11 16:47:43 |
| 1354376 | HZGZS | PRJNA972059 | SAMN35066428 | SRP442780 | SRR24893028 | 2023-06-11 16:47:43 |
| 1354376 | HZGZT | PRJNA972060 | SAMN35066452 | SRP442778 | SRR24893030 | 2023-06-11 16:47:43 |
| 1354376 | HZGZU | PRJNA972061 | SAMN35066487 | SRP442779 | SRR24893029 | 2023-06-11 16:47:43 |
| 1354376 | HZGZW | PRJNA972062 | SAMN35066493 | SRP442777 | SRR24893031 | 2023-06-11 16:47:43 |
| 1354376 | HZGZX | PRJNA972063 | SAMN35066447 | SRP442776 | SRR24893032 | 2023-06-11 16:47:43 |
| 1354377 | HZGZY | PRJNA972064 | SAMN35066484 | SRP442765 | SRR24893011 | 2023-06-11 16:39:15 |
| 1354377 | HZGZZ | PRJNA972065 | SAMN35066466 | SRP442764 | SRR24893012 | 2023-06-11 16:39:15 |
| 1354377 | HZHAA | PRJNA972066 | SAMN35066439 | SRP442760 | SRR24893015 | 2023-06-11 16:39:15 |
| 1354377 | HZHAB | PRJNA972067 | SAMN35066429 | SRP442762 | SRR24893014 | 2023-06-11 16:39:15 |
| 1354377 | HZHAC | PRJNA972068 | SAMN35066494 | SRP442763 | SRR24893013 | 2023-06-11 16:39:15 |
| 1354377 | HZHAG | PRJNA972069 | SAMN35066433 | SRP442759 | SRR24893016 | 2023-06-11 16:39:15 |
| 1354378 | HZHAH | PRJNA972071 | SAMN35066473 | SRP442724 | SRR24892177 | 2023-06-11 16:16:42 |
| 1354378 | HZHAN | PRJNA972072 | SAMN35066409 | SRP442722 | SRR24892179 | 2023-06-11 16:16:42 |
| 1354378 | HZHAO | PRJNA972073 | SAMN35066474 | SRP442723 | SRR24892178 | 2023-06-11 16:16:42 |
| 1354378 | HZHAP | PRJNA972074 | SAMN35066419 | SRP442726 | SRR24892175 | 2023-06-11 16:16:42 |
| 1354378 | HZHAS | PRJNA972075 | SAMN35066503 | SRP442727 | SRR24892174 | 2023-06-11 16:16:42 |
| 1354378 | HZHAT | PRJNA972076 | SAMN35066463 | SRP442725 | SRR24892176 | 2023-06-11 16:16:42 |
| 1354379 | HZHAU | PRJNA972077 | SAMN35066457 | SRP442743 | SRR24892997 | 2023-06-11 16:38:53 |
| 1354379 | HZHAW | PRJNA972078 | SAMN35066416 | SRP442744 | SRR24892996 | 2023-06-11 16:38:53 |
| 1354379 | HZHAX | PRJNA972079 | SAMN35066430 | SRP442745 | SRR24892995 | 2023-06-11 16:38:53 |
| 1354379 | HZHAY | PRJNA972080 | SAMN35066468 | SRP442746 | SRR24892994 | 2023-06-11 16:38:53 |
| 1354379 | HZHAZ | PRJNA972081 | SAMN35066505 | SRP442747 | SRR24892993 | 2023-06-11 16:38:53 |
| 1354379 | HZHBA | PRJNA972082 | SAMN35066420 | SRP442742 | SRR24892998 | 2023-06-11 16:38:53 |
| 1354380 | HZHBB | PRJNA972083 | SAMN35066404 | SRP442740 | SRR24892987 | 2023-06-11 16:39:28 |
| 1354380 | HZHBC | PRJNA972084 | SAMN35066434 | SRP442741 | SRR24892988 | 2023-06-11 16:39:28 |
| 1354380 | HZHBG | PRJNA972085 | SAMN35066417 | SRP442739 | SRR24892989 | 2023-06-11 16:39:28 |
| 1354380 | HZHBH | PRJNA972086 | SAMN35066422 | SRP442736 | SRR24892990 | 2023-06-11 16:39:28 |
| 1354380 | HZHBN | PRJNA972009 | SAMN35066437 | SRP442738 | SRR24892991 | 2023-06-11 16:39:28 |
| 1354380 | HZHBO | PRJNA972010 | SAMN35066499 | SRP442737 | SRR24892992 | 2023-06-11 16:39:28 |
| 1354381 | HZHBP | PRJNA972011 | SAMN35066436 | SRP442752 | SRR24893000 | 2023-06-11 16:46:47 |
| 1354381 | HZHBS | PRJNA972012 | SAMN35066486 | SRP442753 | SRR24892999 | 2023-06-11 16:46:47 |
| 1354381 | HZHBT | PRJNA972013 | SAMN35066500 | SRP442749 | SRR24893002 | 2023-06-11 16:46:47 |
| 1354381 | HZHBU | PRJNA972014 | SAMN35066443 | SRP442751 | SRR24893003 | 2023-06-11 16:46:47 |
| 1354381 | HZHBW | PRJNA972015 | SAMN35066488 | SRP442750 | SRR24893001 | 2023-06-11 16:46:47 |
| 1354381 | HZHBX | PRJNA972016 | SAMN35066410 | SRP442748 | SRR24893004 | 2023-06-11 16:46:47 |
| 1354382 | HZHBY | PRJNA972017 | SAMN35066476 | SRP442768 | SRR24893019 | 2023-06-11 16:38:42 |
| 1354382 | HZHBZ | PRJNA972018 | SAMN35066441 | SRP442770 | SRR24893017 | 2023-06-11 16:38:42 |
| 1354382 | HZHCA | PRJNA972019 | SAMN35066411 | SRP442767 | SRR24893020 | 2023-06-11 16:38:42 |
| 1354382 | HZHCB | PRJNA972020 | SAMN35066485 | SRP442766 | SRR24893021 | 2023-06-11 16:38:42 |
| 1354382 | HZHCC | PRJNA972021 | SAMN35066421 | SRP442769 | SRR24893018 | 2023-06-11 16:38:42 |
| 1354383 | HZHCG | PRJNA972022 | SAMN35066438 | SRP442772 | SRR24893025 | 2023-06-11 17:14:01 |
| 1354383 | HZHCH | PRJNA972023 | SAMN35066475 | SRP442774 | SRR24893023 | 2023-06-11 17:14:01 |
| 1354383 | HZHCN | PRJNA972024 | SAMN35066449 | SRP442771 | SRR24893026 | 2023-06-11 17:14:01 |
| 1354383 | HZHCO | PRJNA972025 | SAMN35066504 | SRP442775 | SRR24893022 | 2023-06-11 17:14:01 |
| 1354383 | HZHCP | PRJNA972026 | SAMN35066462 | SRP442773 | SRR24893024 | 2023-06-11 17:14:0 |

**Supplemental Table 5.** Sample raw and filtered read counts

| Library Name | Raw Reads | Filtered Reads |
| --- | --- | --- |
| HZGZP | 36356482 | 35835980 |
| HZGZS | 64021986 | 62567864 |
| HZGZT | 39893556 | 39358338 |
| HZGZU | 64733806 | 63668178 |
| HZGZW | 75291750 | 73278834 |
| HZGZX | 49028044 | 48453920 |
| HZGZB | 42063742 | 40934728 |
| HZGZC | 63946000 | 61944064 |
| HZGZG | 60110784 | 59522692 |
| HZGZH | 60314034 | 58772452 |
| HZGZN | 51018790 | 50325326 |
| HZGZO | 62425194 | 60617962 |
| HZGZY | 45650338 | 44864526 |
| HZGZZ | 42827352 | 42311188 |
| HZHAA | 46215748 | 45191092 |
| HZHAB | 43171660 | 42620938 |
| HZHAC | 31731726 | 31000840 |
| HZHAG | 49535834 | 47973992 |
| HZHAH | 51549522 | 50946654 |
| HZHAN | 72079986 | 70930308 |
| HZHAO | 47799254 | 47356690 |
| HZHAP | 52442048 | 49592698 |
| HZHAS | 51764020 | 50938202 |
| HZHAT | 55387770 | 50344182 |
| HZHBB | 47852890 | 47194860 |
| HZHBC | 42198512 | 42082904 |
| HZHBG | 72656568 | 72157764 |
| HZHBH | 68576110 | 67166736 |
| HZHBN | 60870270 | 60646210 |
| HZHBO | 64610930 | 63363112 |
| HZHBP | 68307876 | 67337570 |
| HZHBS | 53329198 | 52349752 |
| HZHBT | 43534158 | 42709424 |
| HZHBU | 43075602 | 40703236 |
| HZHBW | 47050642 | 44988738 |
| HZHBX | 50633682 | 48763610 |
| HZHBY | 42654332 | 41902162 |
| HZHBZ | 69647200 | 69002360 |
| HZHCA | 37989808 | 37193176 |
| HZHCB | 49452488 | 48693708 |
| HZHCC | 43114784 | 41534944 |
| HZHCG | 58682442 | 57903410 |
| HZHCH | 41351658 | 40425400 |
| HZHCN | 214601436 | 208096790 |
| HZHCO | 59309864 | 56629082 |
| HZHCP | 188645656 | 180773578 |
| HZHAU | 58637720 | 57889036 |
| HZHAW | 55443470 | 54737262 |
| HZHAX | 46655626 | 45526008 |
| HZHAY | 6251448 | 5448864 |
| HZHAZ | 8083994 | 1688610 |
| HZHBA | 10086490 | 9105258 |
| HZGYU | 81651674 | 79835824 |
| HZGZA | 40559178 | 40144814 |
| HZGYX | 61037212 | 59738782 |
| HZGYY | 62868986 | 61922184 |
| HZGYZ | 35043898 | 34206544 |
| HZGYW | 53404426 | 52939158 |
